# AWC mediated behavioral plasticity in *C. elegans* against *Salmonella* Typhimurium infection

**DOI:** 10.1101/2023.03.28.534663

**Authors:** Swarupa Mallick, Jasmin Pradhan, Vidya Devi Negi

## Abstract

Soil-dwelling nematode *Caenorhabditis elegans* (*C. elegans*) is widely found in close vicinity with different types of microbes, including bacteria, fungus, viruses, etc. However, sensing environmental stress, they often undergo a dormant state called dauer for better survival. Our current study aims to decipher chemosensory responses of worms under *Salmonella* Typhimurium (WT-STM) infection and how bacterial gene modulating worms’ chemosensory system to mediate dauer larvae development. We initially observed the olfactory preference of *C. elegans* toward the pathogenic WT-STM. Although prolonged exposure showed enhanced lawn occupancy of worms in *fepB* mutant *Salmonella* strain with better associative learning response compared to WT-STM counterpart. We also found strong participation of AWC neuron for sensing Δ*fepB* strain and mediating worms’ behavioral plasticity. Overall out study implying a relationship between chemosensory neurons and bacteria emitted signals alter worms’ behavioral plasticity which help us to understand complex scenario of host-pathogen interaction.

## 1. Introduction

Food is the ultimate necessity for organisms’ optimum fitness and survival [1]. However, predation or pathogen attack are quite common for animals while searching food in their own niches. *Caenorhabditis elegans* (*C. elegans*) is free-living soil nematode commonly found in complex environments (decaying vegetation, rotten fruits and compost) in a close association with both pathogenic and non-pathogenic microorganisms which build a hostile situation for worm’s optimum survival. However, *C. elegans* have well-developed chemosensory system with 302 neurons that enable the worms to locate food and sense danger or other animals in their natural habitat [2, 3]. Such chemosensory neurons will translate the chemical cues perceived from its environment and initiate neuroendocrine signaling which allow worms to discriminate between edible food and pathogen contaminated food and drive physical avoidance and also lead to developmental decision in worms [4, 5].

*Salmonella* Typhimurium 14028s strain (WT-STM), a Gram-negative, facultative anaerobic bacteria is well known to cause persistent infection in *C. elegans* and leading to their death [6, 7]. Our previous study demonstrated that deletion of *fepB* genes from bacterial genome enable worms to enter into a dauer stage deciphering food providing some signals, which modulate worms normal physiology [8]. In our current study, we are interested to explore the mechanisms of worms’ chemosensory response toward this Δ*fepB* strain and how such chemosensation mediate worms’ developmental decision under Δ*fepB* strain infection condition. We initially found worms strong food preference towards WT-STM instead of Δ*fepB* strain but, showed better learning phenomenon to avoid the mutant *Salmonella* strain. Besides, we observed significant involvement of AWC olfactory neuron for recognizing Δ*fepB* strain which later involved in mediating developmental plasticity in worms’ population against Δ*fepB* strain. Our study elucidating bacteria emitted signal and organism’s physiology can be used to understand complex host-pathogen interaction in the field of *Salmonella* pathogenesis.

## 2. Materials and methods

### 2.1. Bacterial Strains and growth conditions

*Escherichia coli* strain OP50 (*E. coli* OP50) was a kind gift by Dr. Varsha Singh, Molecular Reproduction, Development and Genetics, IISc. Bangalore, India. *E. coli* OP50 was grown in LB broth and agar and maintained at 37 °C. Wild type *Salmonella* Typhimurium 14028 (WT-STM) was a kind gift from Prof. Dipshikha Chakravortty, Microbiology and Cell Biology, IISc. Bangalore, India. *E. coli* OP50 (RFP), WT-STM (RFP), Δ*fepB* (RFP) and Δ*fepB*_c_ were generated in lab and grown in Luria Bertani (LB) broth and agar with specific antibiotics, e.g., Kanamycin (50 µg/mL) and Ampicillin (50 µg/mL) [(HiMedia laboratory, Mumbai,India)] and kept at 37 °C.

### 2.2. *Caenorhabditis elegans* and their maintenance

*Caenorhabditis elegans* N2 Bristol Wild Type, CX4 [odr-7(ky4) X], AWC ablated worm strains were a kind gift from Dr. Varsha Singh, MRDG, IISc Bangalore’; CX2205 [odr-3(n2150) V] mutant purchased from CGC and FK311 [ceh-36(ks86) X], RB2464 [tax-2(ok3403) I], and VC3113 [tax-4(ok3771) III] were a kind gift from Dr. Kavita Babu, Centre for Neuroscience, IISc Bangalore, India. Worms were cultivated at 22 °C on Nematode Growth Media (NGM) seeded with regular laboratory food, i.e., *E. coli* OP50.

### 2.3. *C. elegans* survival assay

The survival assay was performed as described earlier [8]. In brief 60 L4 worms (N2 and AWC ablated) were transferred to NGM plates seeded with *E. coli* OP50, WT-STM, Δ*fepB*, and Δ*fepB*_*c*_ and kept for 12 hours for infection at 22 °C. Then worms were transferred to the fresh NGM plate seeded with *E. coli* OP50 and counted as day 0, and then each day, worms were transferred to the new *E. coli* OP50 seeded NGM plate and scored for live or dead [9].

### 2.4. Bacterial food choice assay

Overnight bacterial cultures were resuspended in LB broth at O.D. 0.3. Now 25 µl of WT-STM /Δ*fepB* / Δ*fepB*_c_ bacterial suspensions were spotted at one side of 50 mm NGM plate with *E. coli* OP50 on other side and incubated at 37 °C for 12 hours. 50-100 one-day adult (N2) animals were washed 3 times with M9 (Composition for 1 L: KH2PO4-3 g, Na2HPO4-6 g, NaCl-5 g, 1M MgSO4-1 mL) [(HiMedia laboratory, Mumbai, India)] buffer and placed at the center of the assay plates. Animals were allowed to roam for 1-2 hours. Then 5 µl of 1 M sodium azide [(HiMedia laboratory, Mumbai, India)] was added to each bacterial patch to immobilize worms and checked their food preference by using following choice index equation [5].

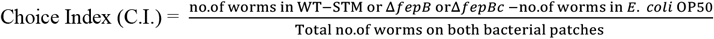

### 2.5. Aversion response assay

50 µl (O.D. 0.3) of WT-STM /Δ*fepB*/ Δ*fepB*_*c*_ bacterial suspensions were spotted middle of the 50 mm NGM plate, and incubated at 37 °C for 12 hours. 100 one-day adult N2 and olfactory neuron defective animals were washed three times with M9 buffer and placed at the center of the assay plates. Aversion response was monitored at different time intervals (12 hours, 24 hours, 36 hours, and 48 hours post-infection). Data was plotted based on their position in bacterial lawn and calculated using following (%) lawn occupancy equation [10].

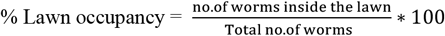

### 2.6. Olfactory aversive learning assay

Around 200 age-synchronized L4 worms (N2 and olfactory neuron defective animals) were transferred to an NGM plate seeded with *E. coli* OP50 (untrained plate), and 100 worms were transferred to four different NGM plates seeded at one corner with 50 µl of *E. coli* OP50 and 200 µl of WT-STM /Δ*fepB*/Δ*fepB*_*c*_ strains to another corner (trained plate). After 48 hours of incubation at 22 °C, 30 worms from trained and untrained plates are transferred to each 50 mm of assay plates containing 50 µl of *E. coli* OP50 and WT-STM /Δ*fepB*/Δ*fepB*_*c*_ strains and observed after 2 hours of incubation. Then 5 µl of 1 M sodium azide [(HiMedia laboratory, Mumbai, India)] was added to each bacterial patch to immobilize worms and checked their food preference and aversive learning behaviour by using the following choice index and learning index equations [11].

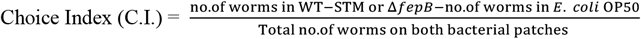

Learning Index (L.I.) = worms in naïve plate – worms in trained plate

### 2.7. Quantification of pharyngeal pumping in *C. elegans*

The 10 L4 age synchronized worms were transferred to NGM plate seeded with *E. coli* OP50, WT-STM, *ΔfepB* and *ΔfepB*_c_. Pharyngeal pumping rate of worms were measured manually under 10X magnification of bright field microscopes and plotted [12, 13].

### 2.8. Quantification of worms’ intestinal lumen width

50 µl (O.D. 0.3) of *E. coli* OP50 and WT-STM/Δ*fepB* bacterial suspensions carrying RFP was spotted middle of the 50 mm NGM plate and incubated at 37 °C for 12 hours. 30 young adult animals were placed inside the bacterial lawn, and at different time intervals (12 to 48 hours post-infection), worms were picked randomly from inside or outside the lawn. Worms were washed 3 times with M9 buffer, mounted on a 2 % agar [(HiMedia laboratory, Mumbai, India)] pad, and visualized under the confocal laser scanning microscope (CLSM) (Leica Microsystems, Wetzlar, Germany). The width of the lumen was analysed using ImageJ software and plotted [10].

### 2.9. Quantitative real-time PCR (qRT-PCR)

RNA from 12 hours to 48 hours infected worms were extracted using TRIzol reagent (Invitrogen, California, USA), as per the manufacturer’s instructions. After the quality check, DNase (Promega, Madison, USA) treatment and control PCR, 2 μg of RNA was used for cDNA synthesis by Go Script TM Reverse Transcription System (Promega, Madison, USA). Then qRT-PCR was performed using Maxima SYBR Green/ROX qPCR master mix (Thermo Scientific, Waltham, USA) with gene-specific primers (Sigma, Bangalore, India) (**Table 1**) in Realplex4 Eppendorf system. Relative expression of the gene was calculated using 2^-ΔΔCT^ method and plotted as fold change [14].

**Table 1.**
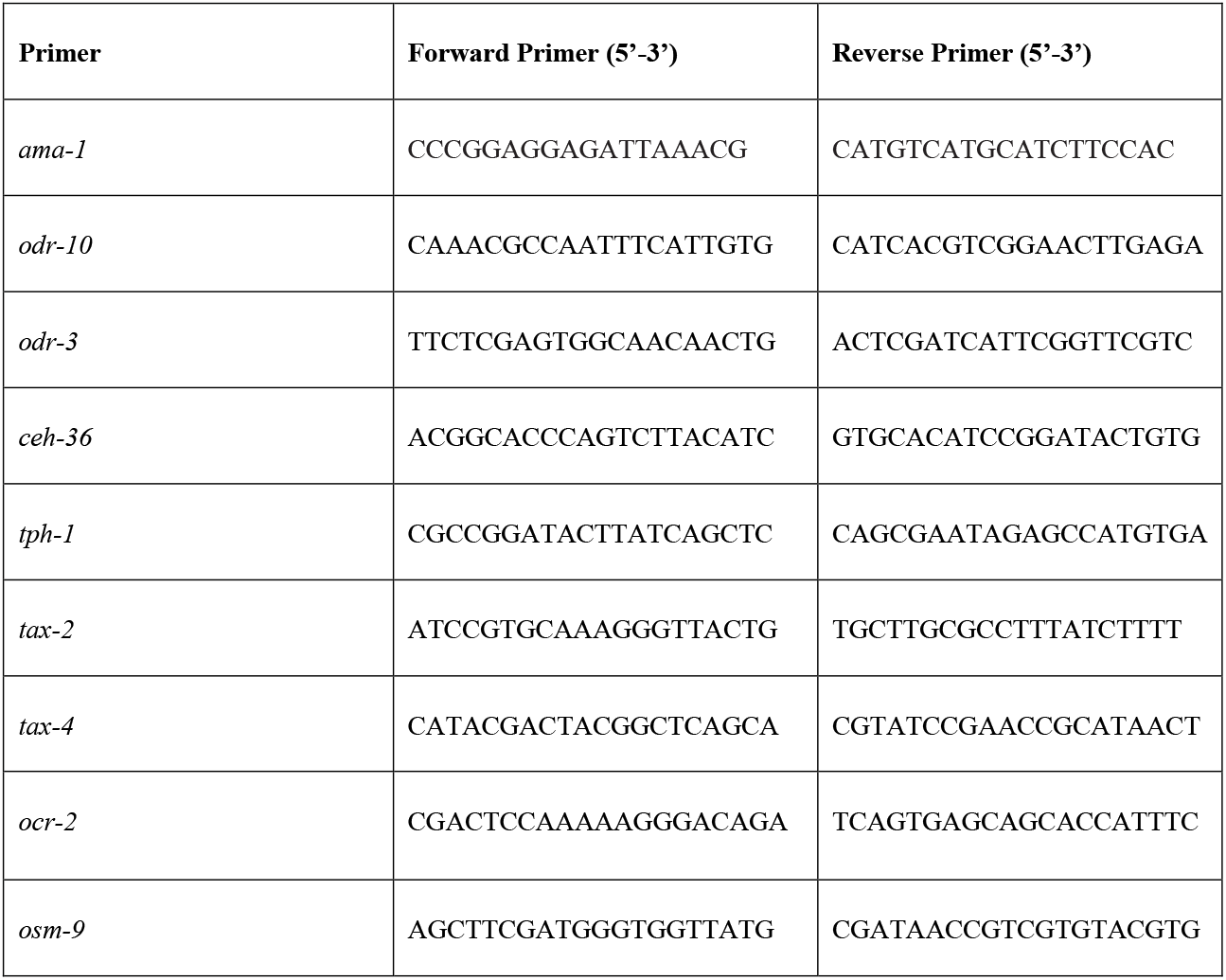
List of primers used in Study.

### 2.10. Statistical Analysis

The Results obtained are represented statistically as mean ± SE. To calculate the statistical significance paired t-test, One-way Anova using Bonferroni’s multiple comparison test and for survival assay log-rank test was performed. To analyze the statistically significant results, we considered a p-value less than 0.05 as statistically significant.

## 3. Results

### 3.1 *Salmonella fepB* mutant infection caused behavioral alteration in *C. elegans*

*C. elegans* can recognize their pathogens and mount certain protective responses like avoidance, resistance or secretion of AMPs, ROS and iNOS for optimum survival under such unfavorable conditions [5, 15]. Here, we were interested to explore the worm’s behavioral responses against *Salmonella* Typhimurium and Δ*fepB* strain. Initially we checked worms’ food preference by performing binary food choice assay and found worms’ strong food preference toward WT-STM over 1-2 hours of exposure (**Fig. 1.A-B**). However, checking lawn aversion behavior of worms against the said pathogens, we observed that in a time dependent manner worms were showing strong lawn aversion against WT-STM but not Δ*fepB* strain (**Fig. 1.C**). Often sensing pathogenic bacteria like *Pseudomonas aeruginosa* (*P. aeruginosa*), *Serratia marcescens* (*S. marcescens*), *Staphylococcus aureus* (*S. aureus*), worms exhibit lawn avoidance behavior, a type of energy sparing-protective response against microbes [16-19]. Together these data implying the early food preference of worms for WT-STM but prolong exposure led to the food preference for Δ*fepB* strain that WT-STM counterpart.

**Fig 1.**
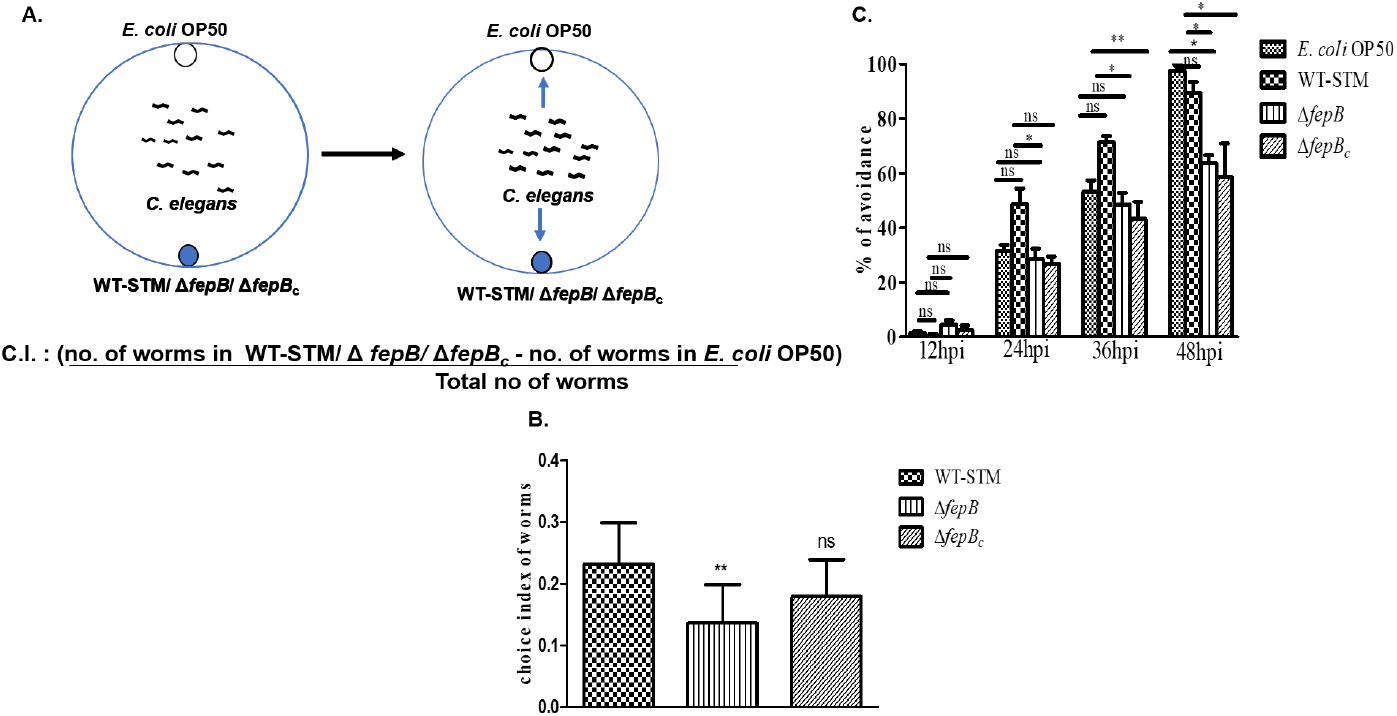
*C. elegans* exhibited altered chemosensation against *fepB* mutant *Salmonella* strain. A. Schematic presentation of binary food choice assay where age-synchronized one-day adult *C. elegans* were placed at the middle of NGM plates seeded with *E. coli* OP50 at one side of the plate and *Salmonella* Typhimurium (WT-STM)/ Δ*fepB/* Δ*fepB*_*c*_ on another side of the plate. Worms were kept under this condition for 1-2 hours and observed their food preference. B. Quantitative analysis of *C. elegans* food preference towards WT-STM/ Δ*fepB/* Δ*fepB*_*c*_ was plotted and shown in the bar. C. Quantitative analysis of *C. elegans* avoidance response was plotted and shown in the bar. The experiments represented three biological replicates with three technical replicates, and the result significance was quantitated as *p*-value < 0.05, ∗, *p*-value < 0.005, ∗∗ and *p*-value < 0.0001, ∗∗∗, ns-non significant. Values expressed as Mean ±SEM.

Often intestinal distension in worms act as an indicator for worms’ aversive response against pathogens. Since, it was reported that, pathogenic bacteria like *P. aeruginosa*, and *E. fecalis* exhibiting aversive response by causing intestinal distension and bloating [13, 16, 20]. Surprisingly in our study we did not observe intestinal distension and change in pharyngeal pumping rate under Δ*fepB* strain infection condition in a time-dependent manner compared to WT-STM (**Fig. 2.A-C**). Overall, decipher an altered behavioral response of worms against pathogens, where worms’ initial food preference was dominated for WT-STM instead of Δ*fepB* although prolong exposure toward Δ*fepB* strain showed worms’ normal uptake of this bacteria with better lawn occupancy.

**Fig 2.**
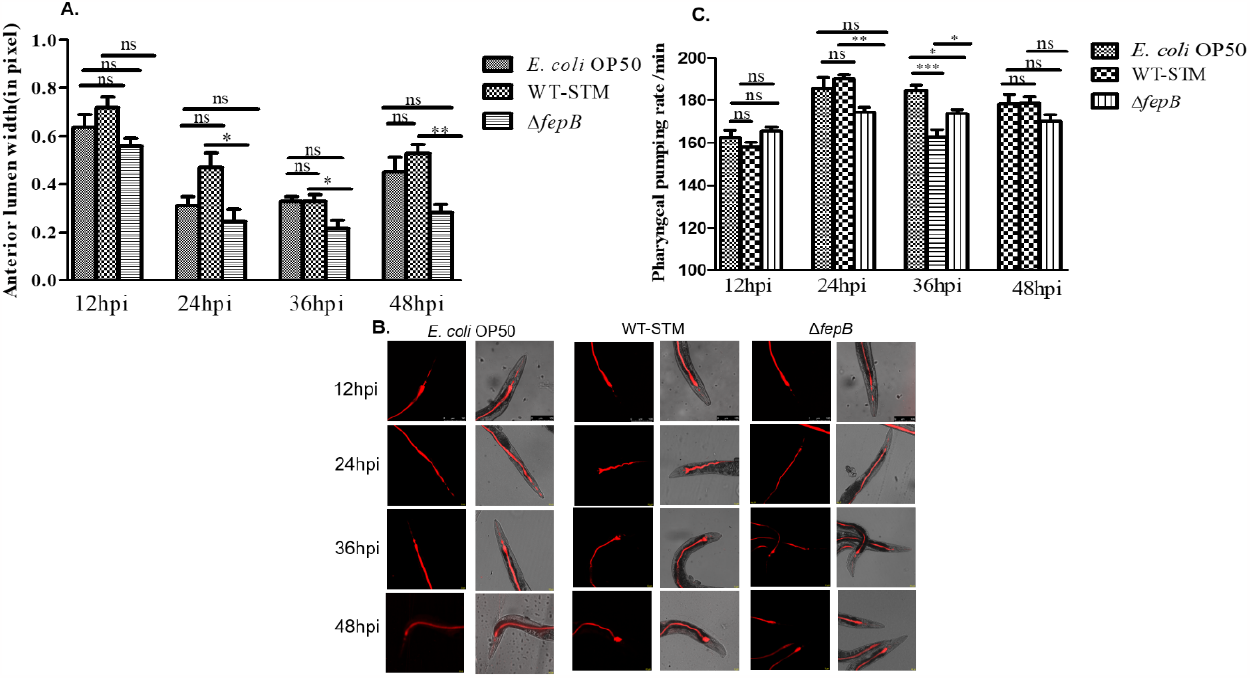
*C. elegans* have no intestinal distention upon Δ*fepB Salmonella* strain infection. Age-synchronized one-day adult *C. elegans* were exposed to RFP plasmid bearing *E. coli* OP50, WT-STM, Δ*fepB*, and checked their anterior lumen width in a time course manner. A. Quantitative analysis of *C*. elegans anterior lumen width was plotted and shown in the bar. B. Confocal microscopy image of worms’ anterior part of the lumen at a different time interval. C. Quantitative analysis of worms’ pharyngeal pumping rate per minute was plotted with respect to 12 hours, 24 hours, 36 hours, and 48 hours post-infection and shown in the bar. The experiments represented three biological replicates with three technical replicates, and the result significance was quantitated as *p*-value < 0.05, ∗, *p*-value < 0.005, ∗∗ and *p*-value < 0.0001, ∗∗∗, ns-non significant. Values expressed as Mean ±SEM. Scale bar 20 µm.

### 3.2. *C. elegans* showed enhanced olfactory function toward Δ*fepB Salmonella* strain

Worms can recognize and remember their pathogen for survival and fitness [21-23]. Continuous exposure to pathogenic bacteria such as *S. enterica, S. marcescene, P. aeruginosa, B. thuringiensis* induce worms to develop aversive learning behavior through the altered feeding preference toward non-pathogenic *E. coli* OP50 [15, 21]. From our initial observation, we found that worms have decreased olfactory preference and aversive responses toward *fepB* mutant *Salmonella* strains in a time-dependent manner compared to WT-STM. Thereafter, we were interested in understanding worms olfactory aversive learning behavior against them. We exposed worms in two different conditions, i.e., in condition one (naive), worms were in *E. coli* OP50 seeded NGM plate, and in condition two (trained), worms were in NGM plate containing *E. coli* OP50 seeded at one side and WT-STM/Δ*fepB*/Δ*fepB*_*c*_ on another side of the plate and incubated for 48 hours in order to acclimatize worms with the odor of these said pathogens (**Fig. 3.A**). Surprisingly we observed that worms trained under Δ*fepB* strain infection condition for 48 hours exhibited increased food preference toward *E. coli* OP50 and also developed associative learning behavior against Δ*fepB* than WT-STM counterpart (**Fig. 3.B-C**). It was reported that *tph-1* gene plays a vital role in the worm’s aversive learning behavior [21]. Thereafter, we also measured the mRNA expression of *tph-1* gene in worms infected by WT-STM and Δ*fepB* strain and surprisingly discovered that gradually with the increase duration of infection, Δ*fepB* infection caused upregulation of *tph-1* expression than WT (**Fig. 3.D**), that strongly implied worms’ enhanced olfaction and better learning behavior against mutant *Salmonella* strain.

**Fig 3.**
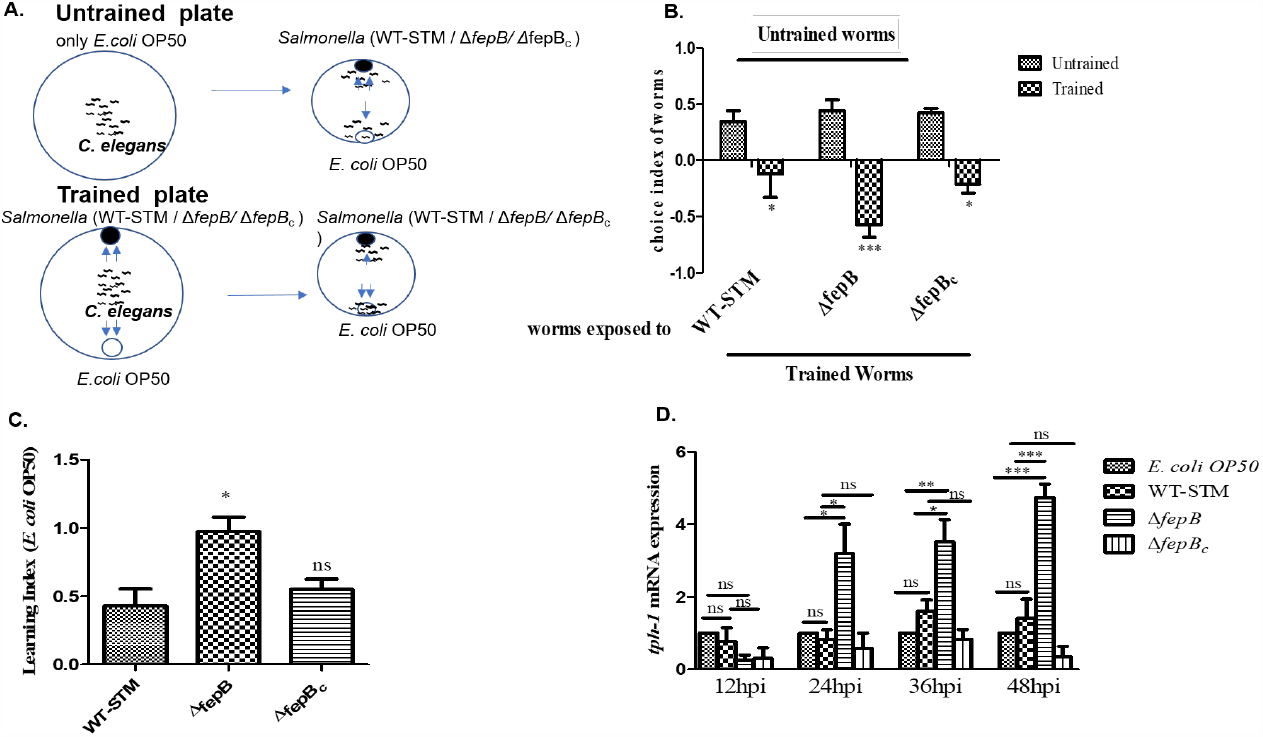
Enhance olfactory function against Δ*fepB Salmonella* strain in *C. elegans*. A. Schematic representation of *C. elegans* olfactory aversive learning behavior where age-synchronized L4 worms were kept in *E. coli* OP50 plate (Naïve plate/untrained plate) and also in NGM plates seeded with *E. coli* OP50 at one side and WT-STM/ Δ*fepB/* Δ*fepB*_*c*_ on another side of the plate (Test plate/trained plate). Worms were kept under this condition for 48 hours, and then worms from both untrained and trained plates were taken, washed 3-4 times, and placed on an assay plate containing *E. coli* OP50 at one side and *Salmonella* Typhimurium (WT-STM)/ Δ*fepB/* Δ*fepB*_*c*_ on another side of the plate. After 1-2 hours worm’s aversive learning behavior was observed. B. Quantitative analysis of worms’ choice index under-trained and untrained conditions were plotted and shown in the bar. Here +1 represents worms’ preference toward WT-STM and mutant strains, -1 represents worm’s preference toward *E. coli* OP50, and 0 represents the equal preference of worms toward *E. coli* OP50 and WT-STM and its mutants. C. Quantitative analysis of worms learning behavior was plotted and shown in the bar. Age-synchronized L4 worms were kept in *E. coli* OP50, plate WT-STM/ Δ*fepB/* Δ*fepB*_*c*_ for 12 hours, 24 hours, 36 hours, and 48 hours. Worms were taken for RNA isolation. After quality and quantity checking, qRT-PCR was performed for *tph-1* genes, *ama-1* was used as the endogenous control, and relative fold changes were calculated using the comparative 2^ΔΔCT^ method. Data represent three biological replicates and three technical replicates, result significance quantitated as *p*-value < 0.05, ∗, *p*-value < 0.005, ∗∗and *p*-value < 0.0 0 01, ∗∗∗, ns-non significant. Values expressed as Mean ±SEM

### 3.3. AWC mediated chemotaxis in *C. elegans* against Δ*fepB* strain

In our study we observed that worms can recognize their pathogens and develop better associative learning to avoid them, particularly Δ*fepB* strain, compared to WT-STM counterparts. However, worms had reduced olfactory preference for Δ*fepB* strain at an early course of the bacterial encounter. This information made us curious to understand the involvement of chemosensory neurons in sensing this particular bacterial strain which was not as pathogenic as WT-STM [8]. Thereafter, we were interested in looking at the participation of olfactory neurons, i.e., AWA, AWB, and AWC, in worms’ olfaction toward the WT-STM and mutant strain along with non-pathogenic *E. coli* OP50 strain as worms can sense volatile odor either attractant or repellent secreted by bacterial strains [4, 5]. From mRNA expression study we observed an upregulation of *odr-10, ceh-36* and *daf-11* at 24 hours post-infection under Δ*fepB* infection condition as compared to WT-STM (**Supplementary Fig. S1**) along with the upregulation of *tax-2* and *tax-4* gene at 24 hours post-infection (**Supplementary Fig. S2**). To confirm the significant involvement of specific neuron in sensing bacterial strain we lawn aversion response of worms having defect in olfactory neurons, i.e., AWA, AWB and AWC and observed that with the increase in time *ceh-36* mutant *C. elegans* remain in Δ*fepB* strain lawn as compare to WT-STM lawn (**Fig. 4.A-D**) indicating *ceh-36* (AWC neuron specific gene) might play an important role in sensing mutant *Salmonella* strain.

**Fig 4.**
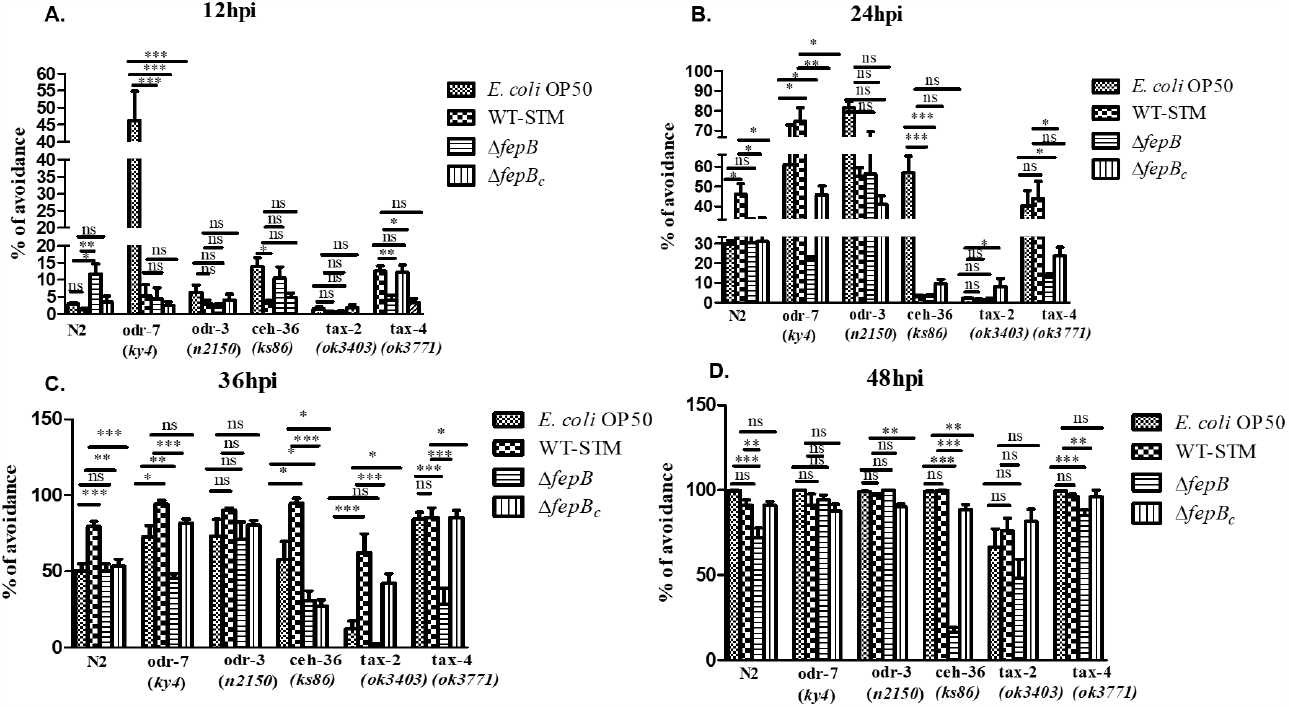
*C. elegans ceh-36* has a role in sensing *fepB* mutant *Salmonella* strain. Age-synchronized one-day adult *C. elegans* (N2, *odr-7, odr-3, ceh-36, tax-2, tax-4*) were exposed to *E. coli* OP50, WT-STM, Δ*fepB*, and Δ*fepB*_*c*_ and observed avoidance behavior in worms in a time-dependent manner. A-D. Quantitative analysis of worms avoidance response was plotted and shown in the bar and was compared with respect to N2 worms. Data represented three biological replicates and three technical replicate, result significance was quantitated as *p*-value < 0.05, ∗, *p*-value < 0.005, ∗∗and *p*-value < 0.0 0 01, ∗∗∗, ns-non significant. Values expressed as Mean ±SEM.

To further confirm the significant involvement of AWC neurons in sensing *fepB* mutant strain, AWC ablated strain was used to check the aversive response. Surprisingly we found that at the later time of infection, i.e., 36 hours and 48 hours post-infection, AWC (-) strain showed significant reduced participation for sensing *fepB* mutant strain as compared to other bacterial strains (**Fig. 5.A-D**). To understand the early cell fate of AWC neuron in sensing Δ*fepB* strain we checked worms’ initial food preferences and olfactory learning behaviour. Surprisingly we found that AWC (-) worms could not sense *Salmonella* Typhimurium, Δ*fepB* strain, and did not even show learning behavior against that pathogen (**Supplementary Fig. S3**). Together our data strongly indicate the role AWC neuron in worms’ chemosensation toward *Salmonella* pathogens, particularly recognizing *fepB* mutant *Salmonella* strain upon continuous infection conditions.

**Fig 5.**
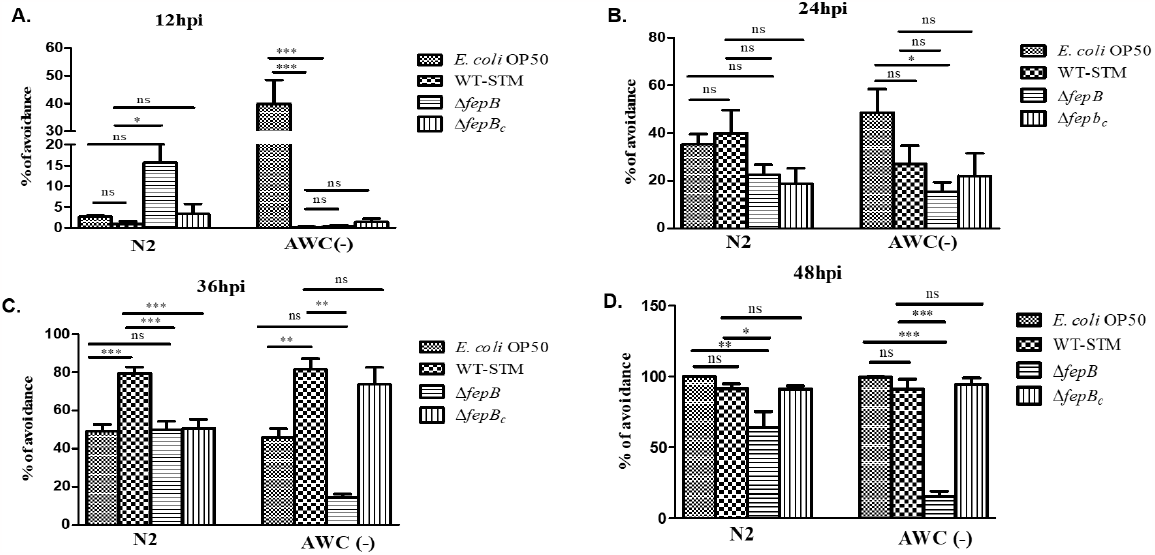
*C. elegans* AWC neurons participate in *fepB* mutant *Salmonella* chemosensation. Age-synchronized one-day adult *C. elegans* [N2, AWC (-)] were exposed to *E. coli* OP50, WT-STM, Δ*fepB*, and Δ*fepB*_*c*_ and observed avoidance behavior in worms in a time-dependent manner. A-D. Quantitative analysis of worms’ avoidance response was plotted and shown in the bar and was compared with respect to N2 worms. Data represent three biological replicated and three technical replicate, result significance was quantitated as *p*-value < 0.05, ∗, *p*-value < 0.005, ∗∗and *p*-value < 0.0 0 01, ∗∗∗, ns-non significant. Values expressed as Mean ±SEM.

### 3.4. AWC neuron involves in behavioral plasticity of worms against Δ*fepB* strain

Often sensing stressful environmental stimuli, i.e., high temperature, pheromone, food limitation, and overcrowding, lead to generating an alternative third larval stage called dauer which can withstand such harsh conditions. Pathogens also act as a stress factor and can modulate worms’ behavioral plasticity [2, 8, 12, 24]. However, the molecular mechanism behind the dauer phenomenon is well studied but how chemosensory neurons participate in raising dauer larvae under pathogen infection needs more study. AWC neuron played a significant role in sensing Δ*fepB* strain. Thereafter, we wanted to understand the involvement of this neuron in modulating worms’ behavioral plasticity. To answer this query, we checked the dauer larvae development under control and infection conditions with olfactory defective mutant worms along with N2 worms. Interestingly we observed reduced number of dauer larvae development by both *ceh-36* and AWC (-) worms in the second generation of worm population and better viability of AWC (-) worms against *fepB* mutant *Salmonella* strain (**Fig. 6. A-C**). Together our study deciphering the major involvement of worm’s AWC neuron in sensing and altered behavioral responses against Δ*fepB Salmonella* strain.

**Fig 6.**
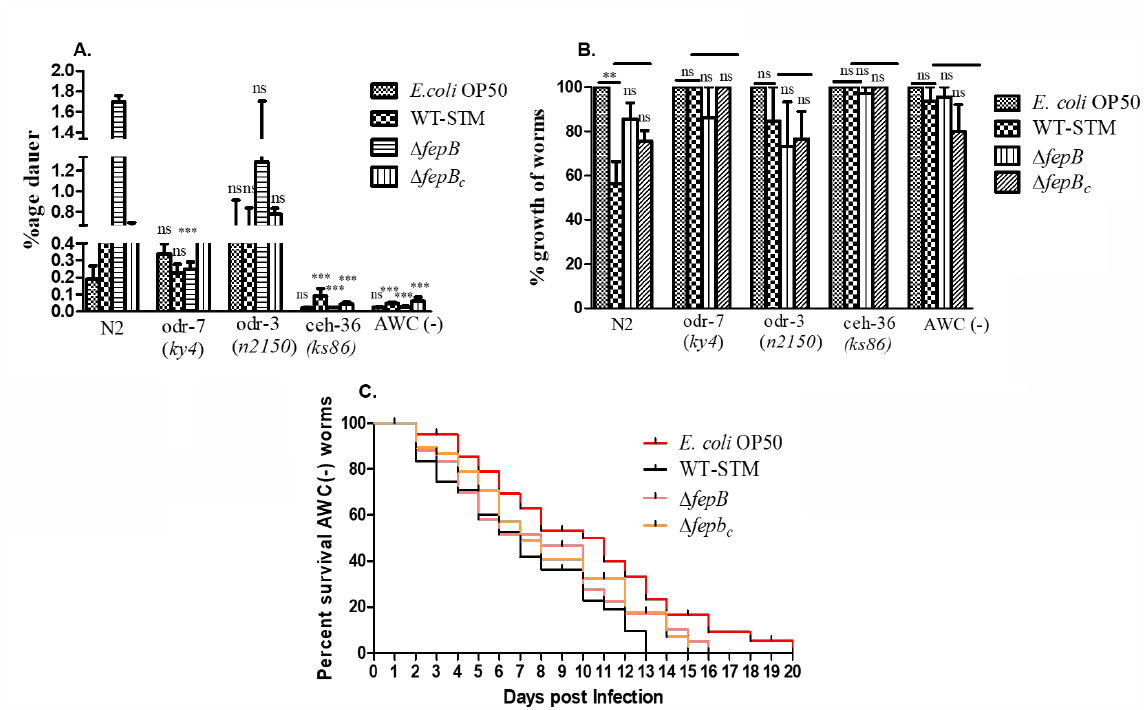
Δ*fepB* strain modulated dauer larva development in *C. elegans* through AWC mediated chemosensation. Age synchronized 10 L4 larvae [N2, *odr-7, odr-3, ceh-36*, AWC (-)] were placed on the NGM plates seeded with non-pathogenic *E. coli* OP50, pathogenic WT-STM, Δ*fepB*, and Δ*fepB*_*c*_ and kept for eight days at 22 °C. On day eight, dauer larvae were isolated using 1% SDS. A. Quantitative analysis of dauer larvae development under control (*E. coli* OP50) and with infection (WT-STM and its mutant strains) conditions were plotted and shown in the bar. Dauer larvae formed under different mutants’ strains were compared with respect to N2 worms. B. AWC (-) worms’ viability was monitored after infecting them for 12 hours with control *E. coli* OP50, WT-STM, Δ*fepB*, and Δ*fepB c* strains (*n* = 30). Data represented three biological replicates and three technical replicate, result significance was quantitated as *p*-value < 0.05, ∗, *p*-value < 0.005, ∗∗and *p*-value < 0.0 0 01, ∗∗∗, ns-non significant. Values expressed as Mean ±SEM.

## 4. Discussion

Animals live in a complex environment that enables them to evolve certain mechanisms to recognize and mount defense responses at molecular and behavioral levels for optimum survival [17, 21]. Bacterivorous nematode *C. elegans* is often found in decaying fruits or other organic matter in close vicinity with commensals and pathogenic microbes. They develop a unique sensorimotor system to navigate their habitat and respond to various environmental cues, including odorant, salt, temperature, pheromone, gases, mechanical stimuli, etc. [15, 17, 25]. In our previous study, we found deletion of *Salmonella* Typhimurium *fepB* gene alters worms’ physiology and develop dauer larva in their second generation [8]. From this initial outcome, we wanted to understand how this Δ*fepB* strain affects worms’ chemosensory system and modulating worms’ behavioral plasticity during infection. We explored worm’s behavioral response by performing binary food choice assay where animal’s initial approach is strongly determined by the bacteria secreted volatile odor [5]. Here we observed worms’ strong olfactory preference toward WT-STM (**Fig. 1.A-B**) but an altered olfactory preference against *Salmonella* strains implying prolong exposure of worms to infection could modulate their behavioral responses (**Fig. 1.C**). However, under Δ*fepB* strain infection condition, we did not observe any intestinal distension as observed in worms under WT-STM infection condition (**Fig. 2.A-B**) and any alteration in the worms’ pharyngeal pumping rate (**Fig. 2.C**). Pathogenic bacteria, i.e., *P. aeruginosa* PA14, *S. marcescens*; fungus *F. oxysporium* infection led to slow death either by secreted secondary metabolite or colonizing the worm’s intestine, causing persistent infection over several days. Still, worms showed aversive learning behavior against those pathogens after worms’ re-exposure to the same pathogen [17, 21, 25, 26]. After giving 48 hours of training to the worms to non-pathogenic *E. coli* OP50 and pathogenic WT-STM and its mutant strain, worms showed an increased associative learning behavior against Δ*fepB* strain than WT-STM counterpart in worms (**Fig. 3.A-C**). Besides, we observed an upregulation of *tph-1* gene (required to synthesize serotonin in worm’s ADF neuron that provides olfactory aversive learning behavior in worms [17, 21, 22]) in a time-dependent manner against Δ*fepB* strain than wild-type *Salmonella* (**Fig. 3.D**) implying worms’ ability to judge food quality, leave the bacterial lawn, and develop an associative learning behavioral by taking their non-pathogenic diet. This finding led us to further explore the involvement of worms’ certain chemosensory neurons for recognizing these bacterial strains.

Worms possess a well-developed chemosensory neuron to sense pathogen-secreted odors that are either attractive or repulsive [5, 16, 27, 28]. There are three olfactory chemosensory neurons located in head region of the worms mostly participate in sensing volatile odors present in their vicinity. Among them AWA and AWC neurons can sense volatile attractive odor whereas AWB neurons can only sense volatile repellent odor [4]. We checked the mRNA expression of these neuron specific genes, i.e., *odr-10, odr-3, ceh-36*. The *odr-10* encode for a G-protein coupled receptor, is expressed exclusively in AWA neuron, and *odr-3* encode for the G protein α subunit required for worms’ normal chemotaxis [4, 29]. Mutation in the *odr-3* gene exhibited defective responses mediated by AWA, AWB, AWC, and ASH neurons in worms. [4, 16, 30, 31]]. *ceh-36* [CEH-36 is a transcription factor playing an essential role in the terminal differentiation of AWC neuron, and mutation to this gene lead to defective responses towards AWC sensed odorant [32]]. Besides, we also measured mRNA expression of the *daf-11* gene since AWC neuron utilized a major signal transduction pathway through the DAF-11 guanylyl cyclase pathway and mutation in *daf-11* gene in animals exhibiting defective responses toward AWC sensed odorant [4, 33, 34]. Mostly chemosensory neurons use cyclic nucleotide-gated ion channels (encoded by *tax-2/tax-4* genes) or TRP (transient receptor potential) channel (encoded by *osm-9/ocr-2* genes) as a secondary sensory transduction pathway [4]. We discovered an upregulation of *odr-10, ceh-36, daf-11* and both *tax-2* and *tax-4* genes at 24 hours post-infection against Δ*fepB* strain infection (**Supplementary Fig. S1, Fig. S2**). We also observed *ceh-36* mutant worms failed to respond to the *fepB* mutant *Salmonella* strain even at 48 hours post-infection than other mutant worms strongly indicating the role of AWC neuron participation for sensing *Salmonella* mutant strain (**Fig. 4.A-D**). to further confirmed the role of AWC neuron, we exposed AWC ablated worms to *E. coli* OP50, WT-STM, Δ*fepB*, and Δ*fepB*_*c*_ strain and observed that AWC (-) worms did not show aversive response against Δ*fepB* at later time of bacterial infection stating strong involvement of AWC olfactory neuron for sensing Δ*fepB* strain (**Fig. 5.A-D**). Behavioral and metabolic changes in organisms are the well-known form of plasticity to withstand harsh environmental conditions [2]. Pathogens also act as stress boosters for inducing dauer larvae in worms. Different groups reported that *P. aeruginosa* PAO1, *S*. Typhimurium MST1, a moderately virulent strain, induces dauer larva formation [12, 24]. Bacteria act as food source for *C. elegans* normal development but how these bacteria are sensed by worms and effecting their developmental process need more study. From our previous study we also observed *fepB* mutant *Salmonella* infection raised significant increase in dauer larva number through the activation of TGF-β signaling pathway [8]. Now, to understand the role of olfactory neurons in modulating worms’ behavioral plasticity we exposed olfactory defective mutant *C. elegans* under continuous infection condition for eight days. Here we discovered significant less dauer larvae number in worm population having defect in AWC neurons as compare to other neurons under Δ*fepB* strain infection (**Fig. 6.A-B**) and also exhibited better viability of AWC (-) of worms under Δ*fepB* strain infection condition compare to WT-STM (**Fig. 6.C**). This study indicating a strong involvement of AWC chemosensory in recognizing *Salmonella* strains and also modulating worms’ behavioral plasticity. Although further study is required to understand whether FepB protein (directly or indirectly) or any other bacterial secretory molecules are involved in altering worm’s behavioral plasticity during the mutant infection that is kept on check in WT-STM. This study will help us to illustrate our knowledge in the field of *Salmonella* pathogenesis and host response, where we speculate the negative regulation of the pathogenesis by *fepB* gene.

## Acknowledgements

Prof. D. Chakravortty, Microbiology and Cell Biology Department, Indian Institute of Science [IISc] Bangalore, India, is acknowledged for providing the *Salmonella* Typhimurium. We are thankful to Canorhabditis Genetic Center (CGC) (University of Minnesota, Twin Cities, USA), Dr Varsha Singh, Molecular Reproduction, Development and Genetics Department IISc., Bangalore, and Dr. Kabita Babu, Centre for Neuroscience, IISc Bangalore for providing the N2 and mutant *C. elegans* strains and *E. coli* OP50 strain. VDN acknowledges the intramural financial support received from the Department of Science and Technology (DST), SERB, Govt. of India EMR/2016/001672 and EEQ/2016/000676 from MHRD, Govt. of India and NIT Rourkela for intramural financial support. SM is MHRD and institutional fellowship recipients and acknowledges the same. We deeply acknowledge Prof. S. K. Patra of Life Science Department and Prof. Sisrendu Sekhar Ray of Biotechnology and Medical Engineering Department of NIT Rourkela for providing the system to perform qRT-PCR for the study.

## Author’s contribution

VDN conceived the idea and supervised the study. VDN and SM planned and designed the experiments. SM performed all the experiments, JP helped in few experiments. VDN and SM analyzed the data and wrote the manuscript.

## Supplementary Figures with legends

**Fig. S1.**
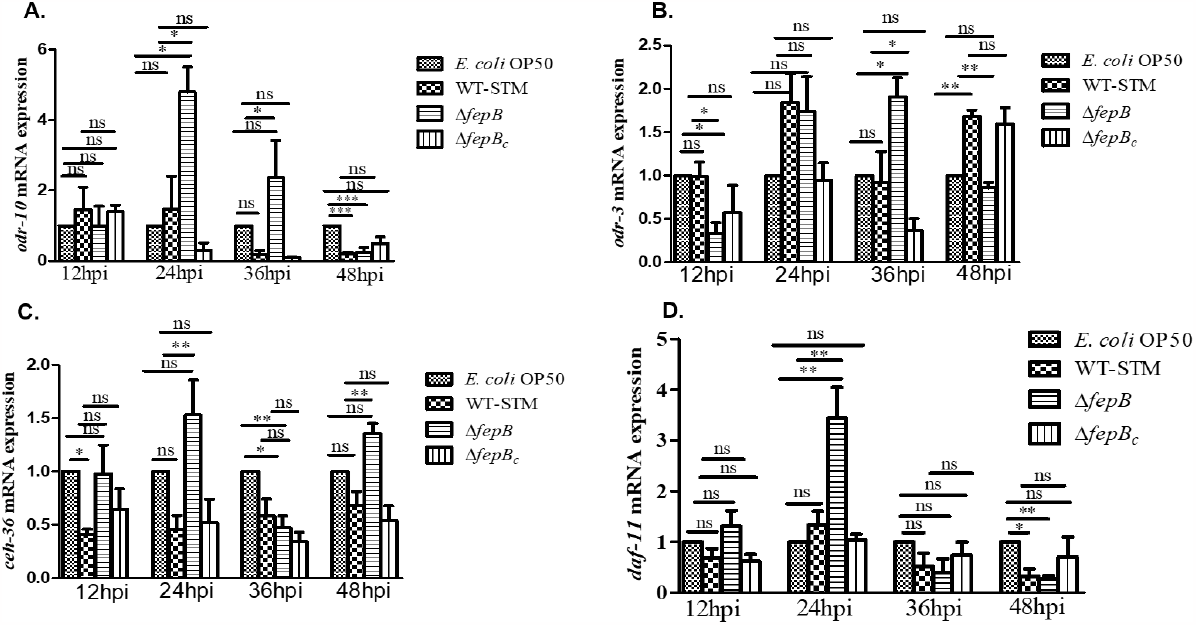
Δ*fepB Salmonella* strain altered olfactory chemosensory response in *C. elegans*. A-D. Age-synchronized L4 worms were kept in *E. coli* OP50, plate WT-STM/ Δ*fepB/* Δ*fepB*_*c*_ for 12 hours, 24 hours, 36 hours, and 48 hours. Worms were taken for RNA isolation, and after quality and quantity checking, qRT-PCR was performed for olfactory neuron-specific genes i.e., *odr-10, odr-3, ceh36, daf-11*; *ama-1* was used as the internal control, and relative fold changes were calculated using the comparative 2^ΔΔCT^ method. Data represented three biological replicates and three technical replicates, result significance was quantitated as *p*-value < 0.05, ∗, *p*-value < 0.005, ∗∗and *p*-value < 0.0 0 01, ∗∗∗, ns-non significant. Values expressed as Mean ±SEM.

**Fig. S2.**
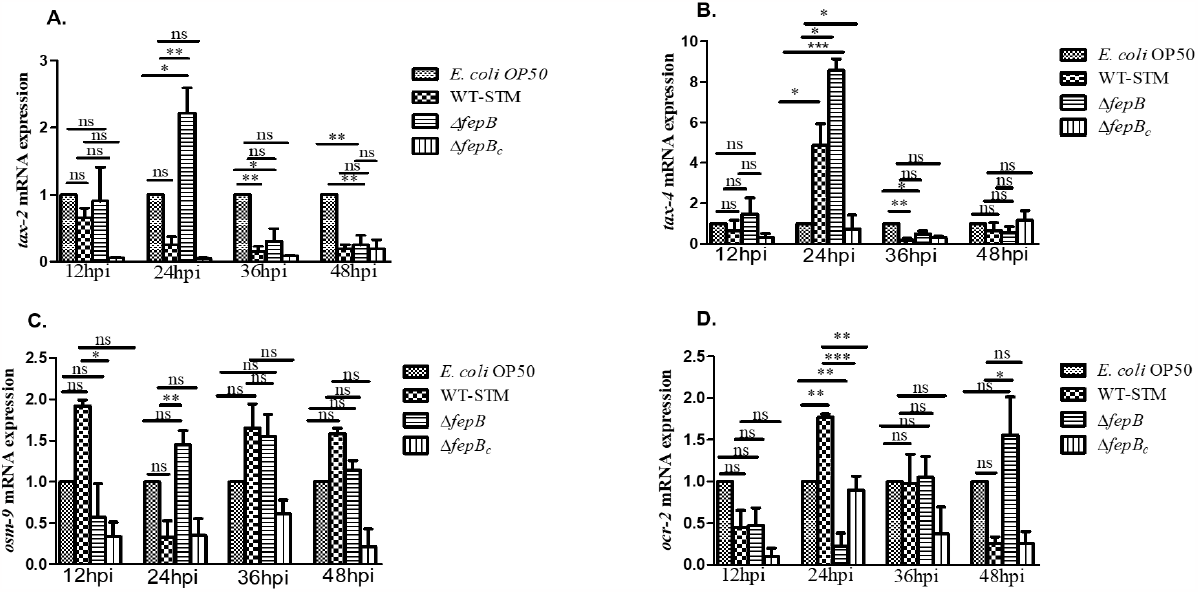
Δ*fepB Salmonella* strain alter olfactory chemosensory response in *C. elegans*. A-D. Age-synchronized L4 worms were kept in *E. coli* OP50, plate WT-STM/ Δ*fepB/* Δ*fepB*_*c*_ for 12 hours, 24 hours, 36 hours, and 48 hours. Worms were taken for RNA isolation, and after quality and quantity checking, qRT-PCR was performed for olfactory neuron-specific genes i.e. *tax-2, tax-4, osm-9, ocr-2*; *ama-1* was used as the endogenous control, and relative fold changes were calculated using the comparative 2^ΔΔCT^ method. Data represented three biological replicates and three technical replicate, result significance was quantitated as *p*-value < 0.05, ∗, *p*-value < 0.005, ∗∗and p-value < 0.0 0 01, ∗∗∗, ns-non significant. Values expressed as Mean ±SEM.

**Fig. S3.**
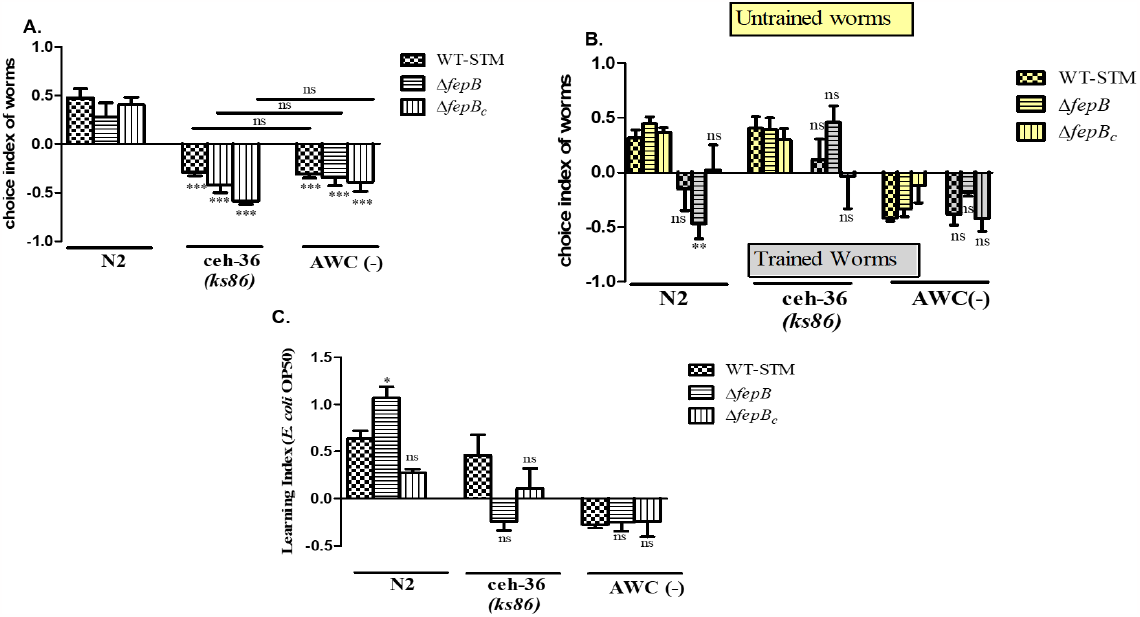
AWC mediated chemotaxis in *C. elegans* against Δ*fepB* strain. Age-synchronized one-day adult *C. elegans* (N2, AWC ablated strain) were placed in the middle of the NGM plate seeded with *E. coli* OP50 at one side of the plate, and *Salmonella* Typhimurium (WT-STM)/ Δ*fepB/* Δ*fepB*_*c*_ on another side of the plate. Worms were kept under this condition for 1-2 hours and observed their food preference. A. Quantitative analysis of worms’ food choices was plotted and shown in the bar. Age-synchronized L4 worms were kept in an *E. coli* OP50 plate (Naïve plate/untrained plate) and also in an NGM plate seeded with *E. coli* OP50 at one side and *Salmonella* Typhimurium (WT-STM)/ Δ*fepB/* Δ*fepB*_*c*_ on another side of the plate(Test plate/trained plate). Worms were kept under this condition for 48 hours, then worms from both untrained and trained plates were taken, washed 3-4 times, and placed on an assay plate containing *E. coli* OP50 at one side and *Salmonella* Typhimurium (WT-STM)/ Δ*fepB/* Δ*fepB*_*c*_ on another side of the plate. After 1-2 hours worm’s learning behavior was observed. B. Quantitative analysis of worm’s choice index under-trained and untrained conditions was plotted and shown in the bar. C. Quantitative analysis of worms’ learning behavior was plotted and shown in the bar. Data represented three biological replicates and three technical replicate, result significance was quantitated as *p*-value < 0.05, ∗, *p*-value < 0.005, ∗∗and *p*-value < 0.0 0 01, ∗∗∗, ns-non significant. Values expressed as Mean ±SEM.

## References

1. Ackley, B.D.J.C.B., Behavior: Should I Stay or Should I Go? 2019. 29(17): p. R842–R844.

2. Fielenbach, N. and A. Antebi C. elegans dauer formation and the molecular basis of plasticity. Genes &development, 2008. 22(16): p. 2149–2165.

3. Pandey, P., et al., Dauer formation in c. Elegans is modulated through awc and asi-dependent chemosensation. 2021. 8(2).

4. Bargmann, C.I., Chemosensation in C. elegans. WormBook: The online review of C. elegans biology [Internet], 2006.

5. Worthy, S.E., et al., Identification of attractive odorants released by preferred bacterial food found in the natural habitats of C. elegans. PloS one, 2018. 13(7): p. e0201158.

6. Aballay, A., P. Yorgey, and F.M.J.C.B. Ausubel, Salmonella typhimurium proliferates and establishes a persistent infection in the intestine of Caenorhabditis elegans. 2000. 10(23): p. 1539–1542.

7. Labrousse, A., et al., Caenorhabditis elegans is a model host for Salmonella typhimurium. 2000. 10(23): p. 1543–1545.

8. Mallick, S., et al., Salmonella Typhimurium fepB negatively regulates C. elegans behavioral plasticity. Journal of Infection, 2022.

9. Pradhan, D. and V.D. Negi, Repeated in-vitro and in-vivo exposure leads to genetic alteration, adaptations, and hypervirulence in Salmonella. Microbial pathogenesis, 2019. 136: p. 103654.

10. Prakash, D., et al., 1-Undecene from Pseudomonas aeruginosa is an olfactory signal for flight-or-fight response in Caenorhabditis elegans. 2021. 40(13): p. e106938.

11. Nag, P., et al., Interplay of neuronal and non-neuronal genes regulates intestinal DAF-16-mediated immune response during Fusarium infection of Caenorhabditis elegans. 2017. 3(1): p. 1–13.

12. Palominos, M.F., et al., Transgenerational diapause as an avoidance strategy against bacterial pathogens in Caenorhabditis elegans. MBio, 2017. 8(5): p. e01234–17.

13. Singh, J. and A. Aballay, Microbial colonization activates an immune fight-and-flight response via neuroendocrine signaling. Developmental cell, 2019. 49(1): p. 89–99. e4.

14. Zhang, J.D., M. Ruschhaupt, and R. Biczok, ddCt method for qRT–PCR data analysis. 2013, Citeseer.

15. Khan, F., S. Jain, and S.F. Oloketuyi, Bacteria and bacterial products: Foe and friends to Caenorhabditis elegans. Microbiological research, 2018. 215: p. 102–113.

16. Prakash, D., et al., 1-Undecene from Pseudomonas aeruginosa is an olfactory signal for flight-or-fight response in Caenorhabditis elegans. The EMBO journal, 2021: p. e106938.

17. Meisel, J.D. and D.H. Kim, Behavioral avoidance of pathogenic bacteria by Caenorhabditis elegans. Trends in immunology, 2014. 35(10): p. 465–470.

18. Zhang, R. and A. Hou, Host-microbe interactions in Caenorhabditis elegans. International Scholarly Research Notices, 2013. 2013.

19. Pradel, E., et al., Detection and avoidance of a natural product from the pathogenic bacterium Serratia marcescens by Caenorhabditis elegans. 2007. 104(7): p. 2295–2300.

20. Filipowicz, A., J. Lalsiamthara, and A. Aballay, TRPM channels mediate learned pathogen avoidance following intestinal distention. Elife, 2021. 10.

21. Zhang, Y., H. Lu, and C.I. Bargmann, Pathogenic bacteria induce aversive olfactory learning in Caenorhabditis elegans. Nature, 2005. 438(7065): p. 179–184.

22. Sasakura, H. and I. Mori, Behavioral plasticity, learning, and memory in C. elegans. Current opinion in neurobiology, 2013. 23(1): p. 92–99.

23. Chiang, Y.-C., C.-P. Liao, and C.-L. Pan, A serotonergic circuit regulates aversive associative learning under mitochondrial stress in C. elegans. Proceedings of the National Academy of Sciences, 2022. 119(11): p. e2115533119.

24. Khanna, A., et al., A genome-wide screen of bacterial mutants that enhance dauer formation in C. elegans. Scientific reports, 2016. 6: p. 38764.

25. Liu, H. and Y. Zhang, What can a worm learn in a bacteria-rich habitat? Journal of Neurogenetics, 2020. 34(3-4): p. 369–377.

26. Nag, P., et al., Interplay of neuronal and non-neuronal genes regulates intestinal DAF-16-mediated immune response during Fusarium infection of Caenorhabditis elegans. Cell death discovery, 2017. 3: p. 17073.

27. Wibisono, P. and J. Sun, Neuro-immune communication in C. elegans defense against pathogen infection. Current Research in Immunology, 2021.

28. Pradel, E., et al., Detection and avoidance of a natural product from the pathogenic bacterium Serratia marcescens by Caenorhabditis elegans. Proceedings of the National Academy of Sciences, 2007. 104(7): p. 2295–2300.

29. Ferkey, D.M., P. Sengupta, and N.D. L’Etoile, Chemosensory signal transduction in Caenorhabditis elegans. Genetics, 2021. 217(3): p. iyab004.

30. Sagasti, A., et al., Alternative olfactory neuron fates are specified by the LIM homeobox gene lim-4. Genes &development, 1999. 13(14): p. 1794–1806.

31. Troemel, E.R., Chemosensory signaling in C. elegans. Bioessays, 1999. 21(12): p. 1011–1020.

32. Pandey, P., et al., Dauer formation in c. Elegans is modulated through awc and asi-dependent chemosensation. Eneuro, 2021. 8(2).

33. Hart, A.C. and M.Y. Chao, From odors to behaviors in Caenorhabditis elegans. The neurobiology of olfaction, 2010.

34. Birnby, D.A., et al., A transmembrane guanylyl cyclase (DAF-11) and Hsp90 (DAF-21) regulate a common set of chemosensory behaviors in Caenorhabditis elegans. Genetics, 2000. 155(1): p. 85–104.

